# Combined fluorescent seed selection and multiplex CRISPR/Cas9 assembly for fast generation of multiple Arabidopsis mutants

**DOI:** 10.1101/2021.05.20.444986

**Authors:** Robertas Ursache, Satoshi Fujita, Valérie Dénervaud Tendon, Niko Geldner

**Affiliations:** Department of Plant Molecular Biology, University of Lausanne, 1015 Lausanne, Switzerland

## Abstract

Multiplex CRISPR-Cas9-based genome editing is an efficient method for targeted disruption of gene function in plants. Use of CRISPR-Cas9 has increased rapidly in recent years and is becoming a routine method for generating single and higher order Arabidopsis mutants. To facilitate rapid and efficient use of CRISPR/Cas9 for Arabidopsis research, we developed a CRISPR/Cas9-based toolbox for generating large deletions at multiple genomic loci, using two-color fluorescent seed selection. In our system, up-to eight gRNAs can be routinely introduced into a binary vector carrying either FastRed, FastGreen or FastCyan fluorescent seed selection cassette. Both, FastRed and FastGreen binary vectors, can be co-transformed as a cocktail via floral dip to introduce sixteen gRNAs at the same time. The seeds can be screened either for red or green fluorescence, or for the presence of both colors at the same time. Our approach provides fast and flexible cloning, avoids very big constructs and enables screening different order mutants in the same generation. Importantly, in the second generation after transformation, Cas9 free plants are identified simply by screening the dark, non-fluorescent seeds. Our collection of binary vectors allows to choose between two widely-used promoters to drive Cas enzymes, either the egg cell-specific (*pEC1.2*) or ubiquitous promoter (*PcUBi4-2*). Available enzymes are “classical” Cas9, a recently reported, intron-optimized version or Cpf1 (Cas12a). Finally, we have taken care to introduce convenient restriction sites flanking promoter, Cas9 and fluorescent selection cassette in the final T-DNA vectors, thus allowing straightforward swapping of all three elements for further adaptation and improvement of the system.

## INTRODUCTION

Generating targeted genetic changes in living cells and organisms has historically been a great challenge in many species, including plants. Precise editing and regulation of genomic information is essential for understanding gene function, production of new plant traits and developing new plant breeding strategies. During the past decade, technological breakthroughs have finally enabled plant genome editing (Weinthal et al., 2010; Curtin et al., 2012; Malzahn et al., 2017). The final breakthrough was achieved with the discovery of CRISPR/Cas-based systems, a gene editing technology which allows us to knock genes in or out (Atkins and Voytas, 2020; Schindele et al., 2020; Zhang et al., 2020; Huang and Puchta, 2021). Knocking out a gene in plants involves expressing CRISPR/Cas and directing it to a specific genomic locus using a guide RNA. There, Cas protein induces sequence-specific DNA double-strand breaks (DSBs), and the cell’s DNA repair mechanism fixes the cut using non-homologous end joining (NHEJ) or homologous recombination (HDR). NHEJ occurs most often, is highly efficient but inaccurate, and tends to introduce errors in the form of small insertions or deletions that are usually sufficient to knock out the gene. This technique is widely used to produce stable single and multiple mutants in various plant species. It is a very valuable tool in studying gene function, breaking the redundancy in multigene families and developing new plant traits (Liu and Fan, 2014; Ma et al., 2016; Manghwar et al., 2019; Atkins and Voytas, 2020; Gao, 2021; Sukegawa et al., 2021). Although multiple Cas proteins have been tested for gene editing in plants, Cas9 and Cpf1 (now known as Cas12a) are currently most widely used based on the nature of their interference complex and their efficiency (Steinert et al., 2015; Ma et al., 2016; Steinert et al., 2017; Schindele and Puchta, 2019; Wolter and Puchta, 2019; Merker et al., 2020). CRISPR/Cas9 consists of two main components: the Cas nuclease and a guide RNA (gRNA). The gRNA is made up of single guide RNA (sgRNA), a short, 17-20 nucleotide sequence complementary to the target genomic DNA, and a tracr RNA, which serves as a binding scaffold for the Cas nuclease. The sgRNA and Cas protein form a Cas/sgRNA complex which is guided to a specific genomic loci site using Watson-Crick base pairing. This results in the cleavage of target DNA sequences adjacent to PAM (protospacer-adjacent motif), a short, few-nucleotide-long sequence that is crucial for Cas binding (Doudna and Charpentier, 2014; Capdeville et al., 2021). Recently, it was demonstrated that the *Cas9* gene codon usage has a significant impact on Cas9 activity in plants. Moreover, the addition of 13 introns into the *Cas9* coding sequence in combination with two nuclear localization signals (NLS) was reported to lead to higher accumulation of the Cas9 nuclease and significant improvement of editing efficiency (Grützner et al., 2021).

In plants, two types of Agrobacterium-mediated techniques are used to create transgenic lines carrying the CRISPR/ Cas system: *in planta* transformation and callus-based transformation (Clough and Bent, 1998; Newell, 2000; Gelvin, 2003; Zhang et al., 2006; Hwang et al., 2017). The most typical example of *in planta* transformation is floral dip-based, where Arabidopsis egg cells have been suggested to be the target of the T-DNA transferred by Agrobacterium infection (Clough and Bent, 1998; Stuitje et al., 2003; Zhang et al., 2006). In case of callus-based systems, excised or partially disrupted meristems are transformed, subjected to antibiotic or herbicide selection, and then carried through tissue culture to regenerate shoots and roots from the transformed tissues (Schmidt and Willmitzer, 1991; Clarke et al., 1992; Bent, 2000). Arabidopsis is highly amenable to *in planta* transformation, giving it an additional advantage as a simple and efficient model organism, compared to many other plant species where in planta transformation methods have often failed and which have to rely on, work- and time-intensive tissue culture protocols.

In theory, the CRISPR/Cas system should be able to function and induce mutations in egg cells or zygotes. However, several studies have demonstrated that the occurrence of such early, complete mutations in the first generation is rare in Arabidopsis and depends on the strength and specificity of promoters driving the Cas expression. When driven under ubiquitous promoters, such as Cauliflower Mosaic Virus *35S* promoter (*CaMV 35S*) or parsley *PcUBi4-2*, a large number of editing events occur later in development, generating a high degree of mosaic plants in the first generation (T1), indicating that CRISPR/Cas9-induced mutations in Arabidopsis occur mostly after the first embryonic cell division, at various stages of plant development (Fauser et al., 2014; Steinert et al., 2015; Wang et al., 2015; Wolter et al., 2018). The use of more specific promoters to drive the expression of Cas enzymes in Arabidopsis, such as egg cell-specific *pEC1.2*, egg cell and early embryo-specific *pDD45*, or cell division-specific *pYAO* promoter, led to a more efficient generation of non-mosaic, biallelic mutants for multiple target genes in Arabidopsis in the T1 generation (Wang et al., 2015; Yan et al., 2015; Mao et al., 2016; Wolter et al., 2018).

Despite the great potential of CRISPR-Cas9 mutagenesis, the current cloning strategies for assembling multiple gRNA vectors, as well as identification of fully mutated, non-mosaic lines carrying large deletions and the simultaneous mutagenesis of multiple genomic loci still require a significant time investment. To unleash the full potential of CRISPR/Cas9 for plant-based applications, an easy-to-use multiplexed assembly system and a fast and efficient screening method is needed. Here, we developed a toolbox with a straightforward cloning protocol for assembly of multiple gRNAs together into transfer DNA (T-DNA) vectors for Agrobacterium-mediated transformation into Arabidopsis. The assembly is based on an efficient combination of Golden Gate and Gateway cloning methods. The T-DNA transfer vectors contain FastRed, FastGreen or FastCyan fluorescent seed selection markers for a fast seed selection under fluorescent stereomicroscopes. We observed that FastRed and FastGreen cassette containing vectors can be easily co-transformed together as an Agrobacterium cocktail to screen for the presence of both seed selection markers in Arabidopsis seeds. This allows to double the amount of target gRNAs, while avoiding generation of very large constructs. Importantly, it also allows to screen for different combinations of mutants in the same generation. The presence of fluorescent seed markers allows to screen fast and efficiently for successful transformation events, as well for editing events already in T1 generation avoiding the antibiotic-caused stress and plant growth defects. In the T2 generation, Cas9-free plants carrying the desired homozygous events in multiple genes can be easily identified by counterselection of non-fluorescent seeds. Although we use *Cas9* driven from *PcUBi4-2* and *pEC1.2* as examples in this study, we expanded our toolbox by introducing the *asCas9*, *asCpf1* intron-optimized *Cas9* into our T-DNA vectors for fluorescent seed sorting. Finally, to expand the possibilities for future modification of FastRed, FastGreen or FastCyan vectors, we introduced convenient restriction sites flanking promoter, Cas9 and the fluorescent seed selection cassette, to be able to select any promoter, Cas protein or selection cassette of interest. This adds an important element of modularity to our system that is absent in many other current systems.

## RESULTS

### An Efficient Assembly System for Multiplexed CRISPR/Cas9

We sought to design an efficient and easy-to-use CRISPR/Cas9 system for the plant research community. Potential applications of such system would include, but are not limited to: 1) simultaneous targeted mutagenesis at multiple Arabidopsis genomic loci; 2) generation of targeted large deletions; 3) further modification of T-DNA vectors by introducing newly discovered Cas proteins, more efficient promoters to drive Cas expression or desired selection cassette. We therefore developed a system that allows for reliable, routine assembly of multiple gRNAs into a T-DNA destination vector already containing Cas9. The assembly method combines both, Golden Gate assembly and Single Gateway recombination (Fig. 1). Three steps are required for assembly. The first step includes generation of entry clones where sgRNAs are introduced into a vector containing either the *pU6* or *pU3* promoter and gRNA scaffold via a simple oligo annealing technique. This is a single tube reaction and only requires an annealed oligonucleotide pair to serve as the gRNA molecule of choice to be introduced into a convenient *BbsI* restriction site (Fig. 1A and Supplemental Table S1). These gRNA entry clones contain overhangs to enable a one-step Golden Gate assembly. This step of cloning relies on BsaI which belongs to Type IIS restriction enzyme family that cleave outside their respective recognition sequences (Fig. 1B–C and Supplemental Table S2). The set of Golden Gate recipient vectors contain the ccdB counterselection cassette, which is replaced by gRNA expression cassettes via single Golden Gate reaction (Fig. 1B–C). After Golden Gate reaction, the recipient vector will contain all sgRNAs assembled together with corresponding pU6 or *pU3* promoters and flanked by attL1 and attL2 sites for the final single Gateway LR reaction (Fig. 1D). The single Gateway LR reaction in our hands always produced a higher number of colonies, compared to double or triple LR reactions. Therefore, we decided to design all our final T-DNA vectors with attR1 and attR2 sites for robustness and high cloning efficiency. In the final LR reaction, all gRNAs are recombined together in the destination T-DNA vector that contains *Cas9* gene, promoter to drive Cas9 expression and fluorescent seed selection cassette of choice (Fig.1D, Table 1). To this end, we constructed eight Golden Gate entry vectors harboring *pU6* or *pU3*-based cassettes (*pRU41–pRU48*) and six Golden Gate recipient vectors (*pSF463, pSF464, pSF278, pSF279, pSF280 and pRU325*) for testing assembly for up to eight gRNA cassettes into a single vector (Fig.1A–C, Supplemental Tables 1 and 2). We found assembly of all eight gRNAs cassettes via Golden Gate was easily achieved and the efficiency for the Golden Gate assembly into intermediate vectors was generally between 70 - 90% (between 7 and 9 colonies out of 10 had correct assembly of gRNAs). All the colonies generated after the final single Gateway LR reaction were positive. T-DNA vector module contains plasmids carrying *Cas9*, *Cas9i* or *Cpf1* variants that have been previously used in higher plants (Table I and Supplemental Fig. S1A). They are plant codon-optimized *SpCas9, SaCas9, Cas9i* and *AsCpf1* variants (Fauser et al., 2014; Steinert et al., 2015; Tang et al., 2017; Grützner et al., 2021). The expression of the different Cas variants is driven either under *PcUBi4-2* or *pEC1.2* promoter (Steinert et al., 2015; Wang et al., 2015; Wolter et al., 2018). As illustrated in the Figure 1, our assembly of a multiplex CRISPR/Cas9 T-DNA vector takes three steps and requires very basic molecular biology techniques. Importantly, PCR is not used for any cloning step, which reduces the likelihood that mutations will occur within the CRISPR/Cas9 components and obviates the need for control sequencing. Having established the system, we next tested our vector systems for genome editing. The following work mainly focuses on testing vectors expressing *Cas9* and *Cas9i* under *pEC1.2* or *PcUBi4-2* promoters.

**Table 1.** T-DNA vectors for FastRed, FastGreen and FastCyan selection

| Destination Vector | Addgene ID | Gateway adapters | Promoter::Cas cassette | Plant selection | Bacterial selection |
| --- | --- | --- | --- | --- | --- |
| pRU051 | 167677 | attR1-attR2 | *PcUBi4-2::SpCas9* | FastRed | Spectinomycin |
| pRU052 | 167678 | attR1-attR2 | *PcUBi4-2::SpCas9* | FastGreen | Spectinomycin |
| pRU053 | 167679 | attR1-attR2 | *pEC1.2::SpCas9* | FastRed | Spectinomycin |
| pRU054 | 167680 | attR1-attR2 | *pEC1.2::SpCas9* | FastGreen | Spectinomycin |
| pRU321 | 167681 | attR1-attR2 | *PcUBi4-2::SaCas9* | FastRed | Spectinomycin |
| pRU320 | 167682 | attR1-attR2 | *PcUBi4-2::SaCas9* | FastGreen | Spectinomycin |
| pRU322 | 167683 | attR1-attR2 | *pEC1.2::SaCas9* | FastGreen | Spectinomycin |
| pRU055 | 167684 | attR1-attR2 | *pEC1.2::AsCpf1* | FastRed | Spectinomycin |
| pRU205 | 167685 | attR1-attR2 | *PcUBi4-2::AsCpf1* | FastGreen | Spectinomycin |
| pRU319 | 167686 | attR1-attR2 | *pEC1.2::SpCas9* | FastRed | Spectinomycin |
| pRU292 | 167687 | attR1-attR2 | *PcUBi4-2::Cas9i* | FastRed | Spectinomycin |
| pRU293 | 167688 | attR1-attR2 | *pUBi4-2::Cas9i* | FastGreen | Spectinomycin |
| pRU294 | 167689 | attR1-attR2 | *pEC1.2::Cas9i* | FastRed | Spectinomycin |
| pRU323 | 167690 | attR1-attR2 | *PcUBi4-2::Cas9i* | FastCyan | Spectinomycin |
| pRU324 | 167691 | attR1-attR2 | *pEC1.2::Cas9i* | FastCyan | Spectinomycin |
| pRU061 | 167692 | attR1-attR2 | *PcUBi4-2::AsCpf1* | FastRed | Spectinomycin |
| pRU206 | 167693 | attR1-attR2 | *pEC1.2::AsCpf1* | FastGreen | Spectinomycin |
| pRU295 | 167694 | attR1-attR2 | *pEC1.2::Cas9i* | FastGreen | Spectinomycin |

**Figure 1.**
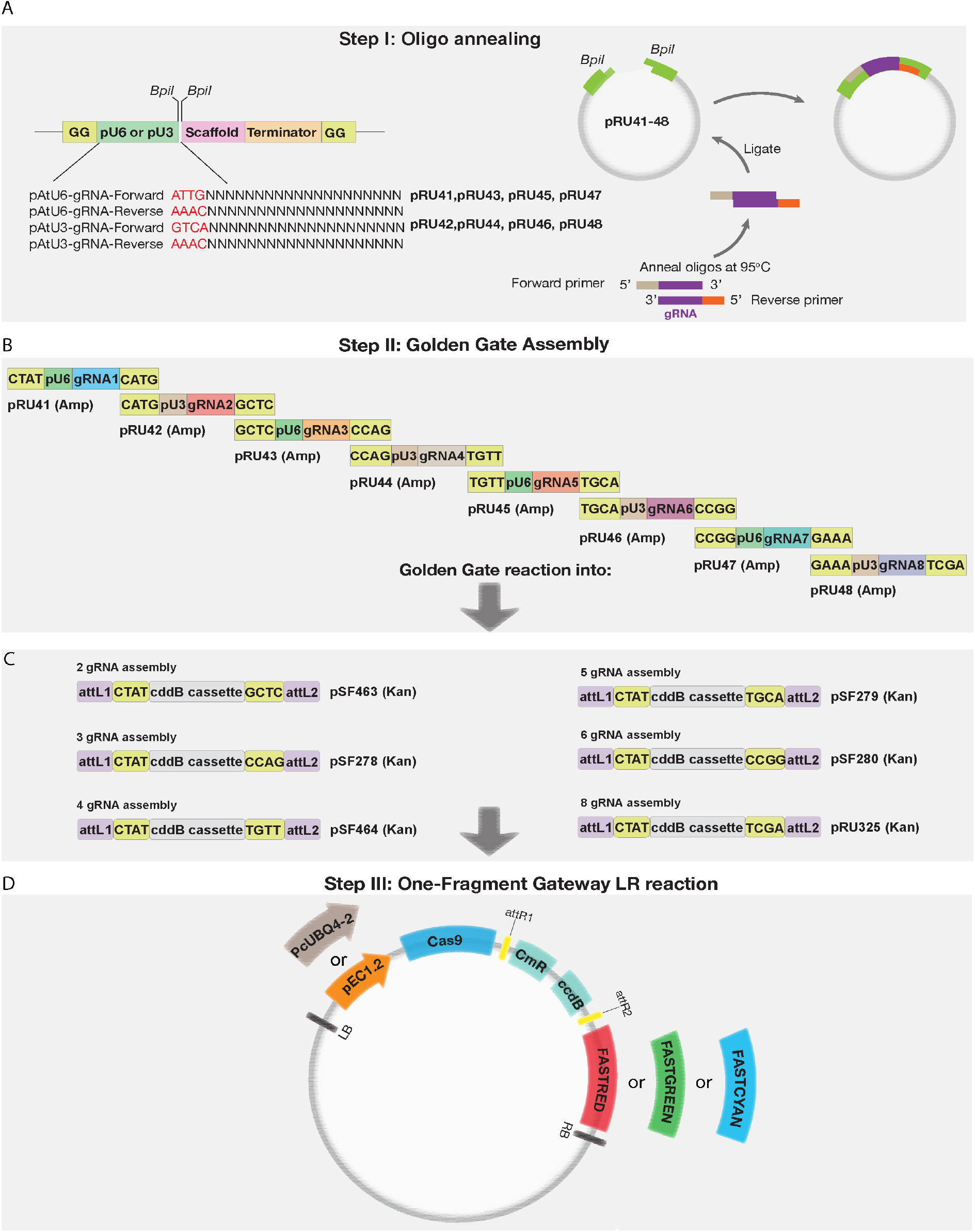
CRISPR/Cas cloning strategy. A, Oligo annealing-based cloning of chosen gRNAs into *pRU41-48* entry vectors containing pU6 and pU3 promoters. B-C, Golden Gate assembly of up to eight gRNAs into corresponding intermediate vectors containing attL1-attL2 for single Gateway LR reaction. D, Final single Gateway assembly into T-DNA vectors containing *PcUBi4-2* or *pEC1.2* promoters driving Cas9 of interest. The vectors contain one of three fluorescent seed selection cassettes, FastRed, FastGreen and FastCyan.

### The game of colors – a cocktail of FastRed and FastGreen vectors for a faster screening of editing events

Into each T-DNA vector carrying the Cas9 cassette we have introduced FastRed or FastGreen or FastCyan fluorescent seed marker for a faster seed sorting (Fig. 1D, Table 1 and Supplemental Table S1A). As described previously, FAST (fluorescence-accumulating seed technology), is based on the expression of OLE1 (OLEOSIN1) translational fusions, under the control of its native pOLE1 promoter, thus accumulating fluorescence on the oil body membranes in the developing seeds of Arabidopsis. Arabidopsis seeds accumulate a large quantity of oil bodies, which are surrounded by phospholipid membranes with embedded proteins within the cell (Huang, 1992). Oleosins are abundant structural proteins embedded in oil body membranes, have an important function in regulating the size of oil bodies and confer freezing tolerance upon seeds (Siloto et al., 2006). Among all oleosins, OLE1 is the most abundant in Arabidopsis seeds (Shimada et al., 2008; Shimada et al., 2010).

We introduced into our CRISPR/Cas constructs a *pOLE1::OLE1* fusion tagged either with *GFP*, *tagRFP* or *mTurquoise2* to be able to screen seeds with red, green or cyan fluorescence simply by using a fluorescent stereomicroscope (Table 1, Figure2). The aim was to reduce the length of time for: 1) identifying the transgenic lines in T1 carrying the desired events; 2) obtaining homozygous, CAS9-free mutants in T2 generation. In addition, we decided to test whether the T-DNA constructs, carrying FastRed or FastGreen, can be combined together via simple Agrobacterium-based co-transformation. Immediately before dipping Arabidopsis flowers, we mixed the two Agrobacterium solutions in equal proportions together into one cocktail.

**Figure 2.**
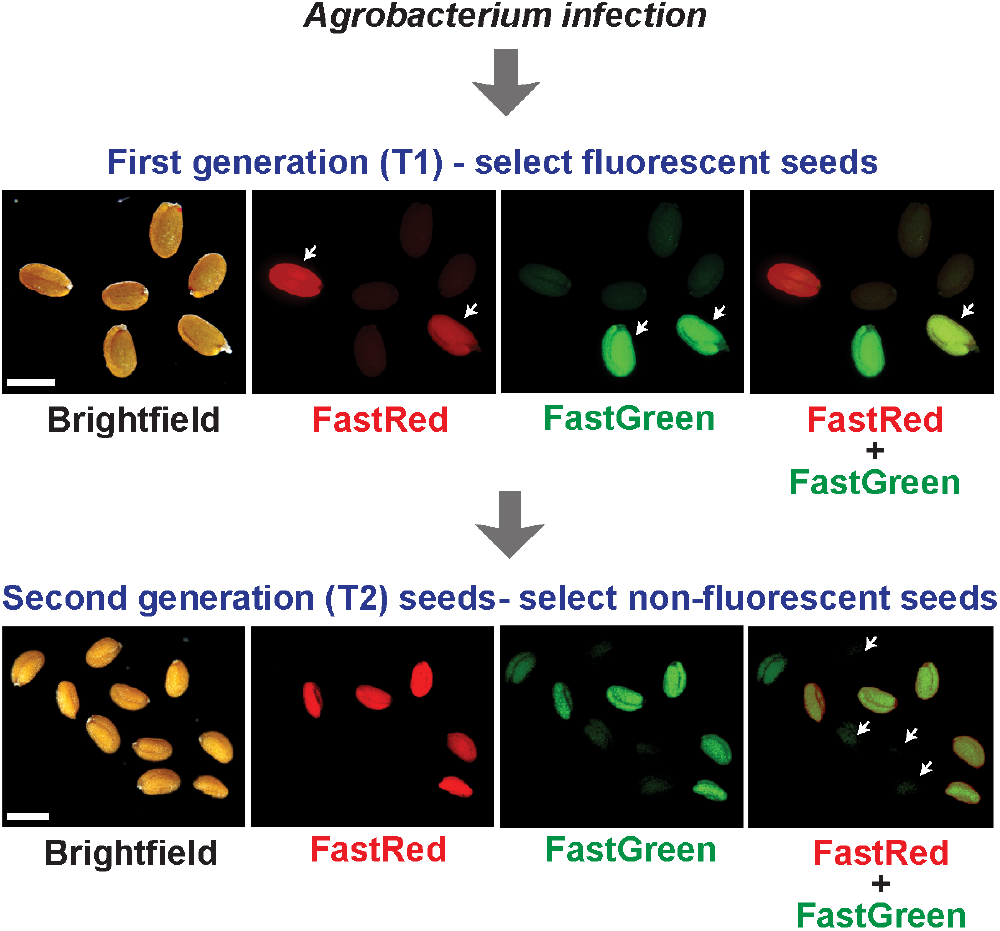
Fluorescent seed selection in T1 and T2 generations after Agrobac-terium-mediated transformation. Green, red or yellow (two color) fluorescent seeds are selected in T1 generation, genotyped for editing events and the dark, Cas9-free, seeds of T2 progenies are screened for stably inherited mutations. Scale bars = 100 μm.

In T1 generation, we were able to screen either for red or green fluorescent seeds, or both colors together (rates of 3-4% single color seeds and 0.5-1% of two-color seeds, based on four independent co-dipping experiments, 16 plants co-dipped in each experiment). Dependent on the filter set available in users’ fluorescent stereomicroscopes, the seeds can be screened for both colors either by switching the filters in-between or by using the long pass filters for green fluorescence to select the yellow color seeds. In T2 generation, we noticed that the majority of two-color seeds lines contained mostly yellow seeds suggesting that, although coming from two separate vectors and independent bacteria, FastRed and FastGreen cassettes containing T-DNAs often enter into a common locus and co-segregate together (Fig. 2 and Supplemental Fig. S1B). Screening of non-fluorescent seeds in T2 (counter-selection) allowed us to obtain Cas9-free homozygous mutants. We used *cuc1cuc2* double mutants as examples to demonstrate the application of our FastRed and FastGreen co-transformation system.

### Simultaneous targeting of different Arabidopsis loci using FastRed and FastGreen strategy

First, we tested our system for simultaneously creating targeted deletions in Arabidopsis *CUC1* and *CUC2* loci. The *CUP-SHAPED COTYLEDON* genes *CUC1* and *CUC2* encode a pair of NAC transcription factors required for shoot meristem initiation. They are functionally redundant and the seedlings of each single mutant show little morphological phenotype while the double mutant completely lacks a shoot meristem and produce completely fused cotyledons (Aida et al., 1999; Aida et al., 2002; Hibara et al., 2003; Hibara et al., 2006). To increase the chance of obtaining different mutant alleles and large deletions, we picked three gRNAs for each, *CUC1* and *CUC2*, located in different exons and distributed along the coding sequence (Fig. 3A). The three gRNAs for each gene were expressed either under the *pU6* or *pU3* promoter as indicated in the cloning setup (Fig. 1A). The T-DNA expression vector for *CUC1* contained FastRed and the vector for *CUC2* – FastGreen selection marker. To compare the efficiencies, we used constructs containing either *pEC1.2* or *PcUBi4-2* promoter driving *Cas9* expression. Both, FastRed and FastGreen constructs, were co-dipped together as an Agrobacterium cocktail into wild-type Col-0 plants to generate seeds with single or both colors. In the first generation after transformation (T1), we selected the fluorescent seeds containing both colors using fluorescent stereomicroscope, germinated them on plates and genotyped. We identified heterozygous (3/19; 15.8 % for *pEC1.2* and 6/22; 22.7% for *PcUBi4-2* construct) and biallelic (3/19; 15.8% efficiency for *pEC1.2* and 3/22; 11.1% for *PcUBi4-2* constructs) double mutants of *cuc1cuc2* (Fig. 3B). We also identified 1/19 (5,3%) and 3/22 (13.6%) chimeric plants in *pEC1.2* and *PcUBi4-2* experiments accordingly. Although the overall number of events was higher in the co-transformation experiments where the *PcUBi4-2* was used to drive *Cas9*, we noticed that more biallelic mutants were found in T1 when *pEC1.2* promoter was used. This is consistent with previous studies where *pEC1.2* was shown to produce biallelic and heritable events already in T1. As expected, the *cuc1cuc2* double mutants exhibited a severe shoot phenotype with fused cotyledons (Fig. 3B). In our hands, the *cuc1cuc2* mutants were not viable and therefore we had to maintain heterozygous T1 lines. To check if the detected mutations in both genes were stably transmitted from heterozygous T1 to T2 generation, we took two heterozygous T1 lines from *pEC1.2::Cas9* and four lines from *PcUBi4-2::Cas9* co-dipping experiment, left them to self-pollinate, checked the segregation and type of mutations in the following T2 generation. The resulting T2 lines segregated and we were able to detect a high rate of seedlings showing double mutant phenotype (Fi. 3C).

**Figure 3.**
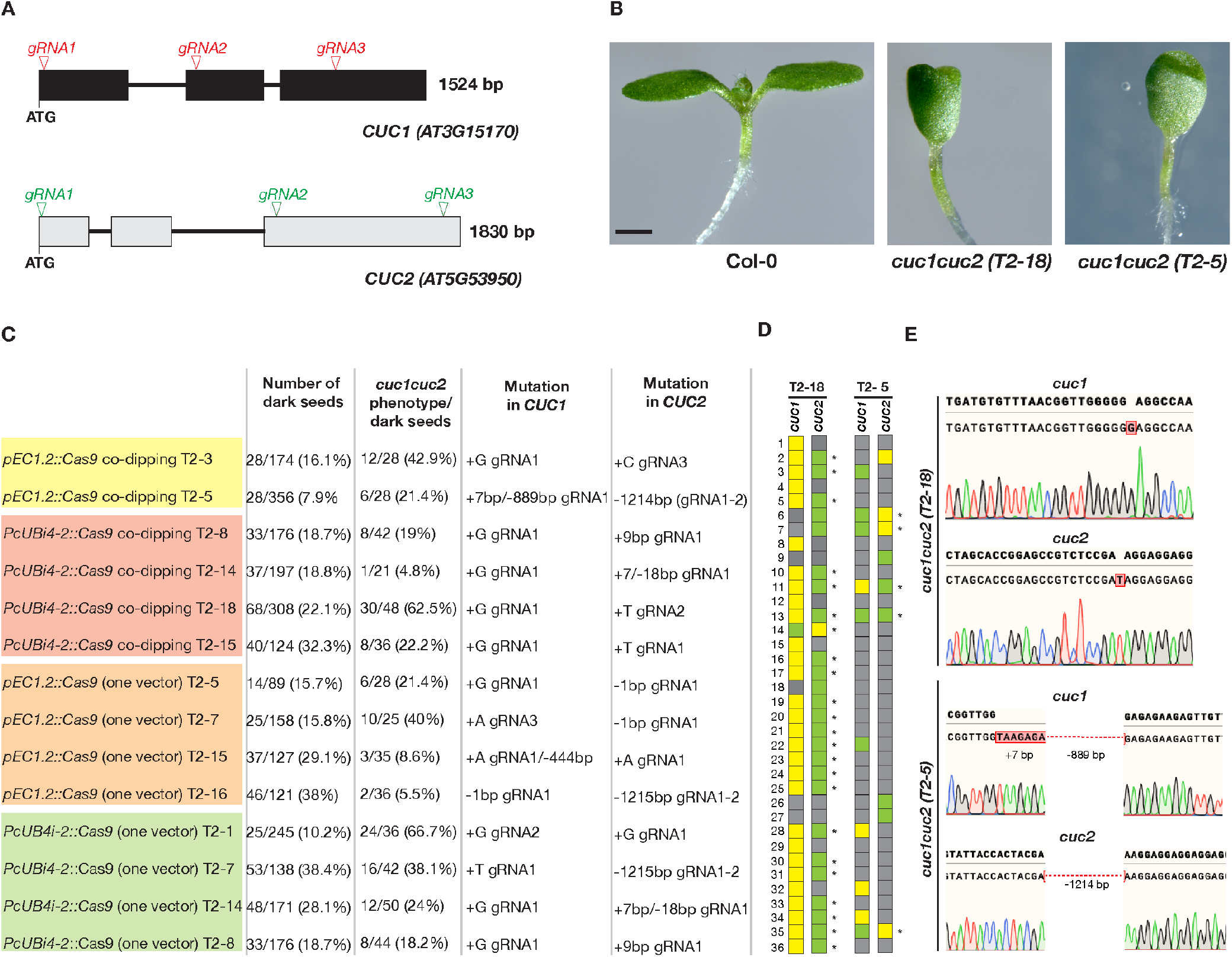
Example of the application of co-dipping strategy. A, Graphical representation of *CUC1* and *CUC2* genes and positions of gRNAs. B, *cuc1cuc2* knockout phenotype in the isolated *pEC1.2::Cas9* (T2-5) and *PcUBi4-2* (T2-18) lines. C, Segregation analysis of the selected T2 lines. Counting the darks seeds and evaluating *cuc1cuc2* phenotype among dark seeds were performed independently. Yellow color stands for homozygous, green for heterozygous and grey color for wild-type individuals. The asterisks indicate the individuals homozygous for one gene and heterozygous for the one. D, Segregation analysis in 36 wild-type looking individuals derived from dark seeds from the selected *pEC1.2::Cas9* (T2-5) and *PcUBi4-2* (T2-18) lines. E, The chromatograms showing the type of mutation in *pEC1.2::Cas9* (T2-5) and *PcUBi4-2::Cas9* (T2-18) lines as identified by sequencing. Scale bar in B = 1mm.

Due to severe cotyledon phenotype, we were not able to maintain homozygous *cuc1cuc2* for next generations. Therefore, in order to get rid of the Cas9-containing T-DNA construct and maintain viable seedlings, we screened only for dark seeds and identified individuals with a homozygous mutation in one of the *CUC* genes and heterozygous for the second one. We sorted 36 dark seeds of two T2 lines, one derived from *pEC1.2::Cas9* (T2-5) and the other one from *PcUBi4-2::Cas9* (T2-18) co-dipping experiments, for more detailed segregation analysis. In case of *pEC1.2::Cas9* (T2-5) line, we detected 5/36 individuals (13.9%) with a heterozygous mutation for one of the *CUC* genes and homozygous for the other gene. In case of *PcUBi4-2::Cas9* (T2-18) line, we found 23/36 individuals (63.9%) heterozygous for one of the *CUC* genes and homozygous for the other gene (Fig. 3D–E). We further verified the absence of Cas9 by PCR using Cas9-specific primers in these lines. Thus, following all these results, our multiplex CRISPR/Cas9 system allows effective expression of at least three gRNAs/gene and the combination of two constructs to target two genomic loci simultaneously. Such strategy also provides flexibility in the screening strategy where generation of different combinations of mutants is required.

To test if we can efficiently target a higher number of gRNAs using the same construct, we cloned three gRNAs for *CUC1* and two gRNAs for *CUC2* into the same vector carrying Cas9 under *pEC1.2* promoter. The resulting vector contained five gRNAs in total. In T1, we could obtain biallelic *cuc1cuc2* (16%; 4/25 for *pEC1.2::Cas9* and 14.3%; 4/28 for *PcUBi4-2::Cas9*), heterozygous (5/25; 25% for *pEC1.2::Cas9* and 7/28; 25% for *PcUBi4-2::Cas9*) with similar ratio compared to the co-transformation strategy. This suggest that stacking up at least five gRNAs into the same vector does not affect the efficiency of Cas9-mediated editing. In T2 generation, we were able to identify a number of segregating lines containing Cas9-free and carrying different types of mutations (Fig. 3C).

Given that the vectors we generated have maximum capacity of eight gRNAs, the number can be doubled to sixteen using co-transformation technique. This provides a powerful tool for generation of high order Arabidopsis mutants. The system described here has been successfully applied to generate several important high order Arabidopsis mutants: quintuple *gelp22/gelp38/ gelp49/gelp51/gelp96* (*gelp^quint^*) mutant lacking the core root endodermis suberin polymerization machinery, nonuple endodermis-specific laccase *lac1;3;5;7;8;9;12;13;16 (9× lac)* mutant and a quadruple *myb41-myb53-myb92-myb93* (*quad-myb*) with mutations in four transcription factors essential for promoting endodermal suberin formation (Rojas-Murcia et al., 2020; Shukla et al., 2021; Ursache et al., 2021).

### Flexible CRISPR/Cas T-DNA vectors for further modification and improvement

Currently, new Cas proteins and their modifications are being discovered and reported at a high rate (Steinert et al., 2015; Schindele and Puchta, 2019; Wolter and Puchta, 2019; Merker et al., 2020; Huang and Puchta, 2021). Some of them more efficient than others, with different binding properties and different PAM sites. Therefore, we decided to modify our vectors in order to enable a fast and easy swapping of the promoter to drive Cas, Cas itself and the plant selection cassette. Our vectors have *KpnI* restriction sites flanking the promoter *pEC1.2*, as well as *SgsI* sites flanking the Cas9 sequence and *HindIII* restriction sites flanking the fluorescent seed selection cassette (Fig. 4A). All these restriction sites allow for a fast and efficient replacement of all three elements in the future. Our vectors could be easily modified for generating tissue-specific CRISPR by replacing *PcUBi4-2* or *pEC1.2* with a tissue-specific promoter of interest, for example.

**Figure 4.**
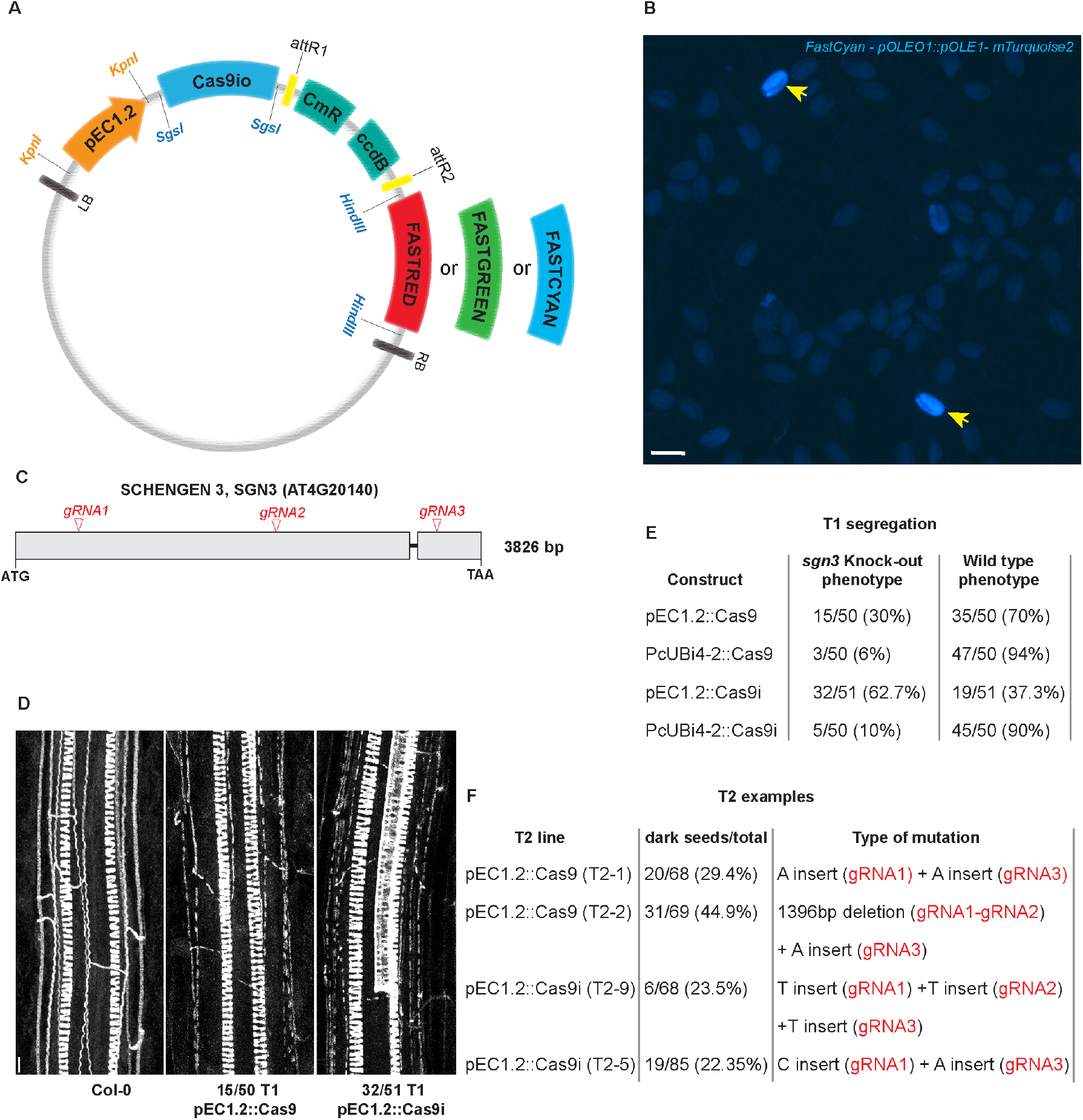
CRISPR/Cas T-DNA vectors for further modification and improvement. A, Schematic representation of T-DNA vector containing *KpnI*, *SgsI* and *HindIII* restrictions sites flanking the promoter, Cas gene and selection cassette, accordingly. B, FastCyan seed selection based on seed-specific *pOLE::OLE1-mTurquoise2* expression. Yellow arrows indicate the positive fluorescent seeds. C, Schematics of *SGN3* gene showing the positions of the chosen gRNAs. D, Basic Fuchsin (gray) staining of lignin-based Casparian strips in the isolated T1 generation biallelic mutants generated using *pEC1.2::Cas9* and *pEC1.2::Cas9i* constructs. E, Segregation analysis in T1 generation in *pEC1.2::Cas9*, *pEC1.2::Cas9i*, *pUBi4-2::Cas9* and *pUBi4-2::Cas9i* lines. Scale bars = 100 μm (B) and in 25 μm (D).

To demonstrate the advantage of our modular cloning system, we decided to replace the Cas9 in the destination vector *pEC1.2::Cas9* with a recently published optimized version of *Cas9*, *Cas9i*, containing 13 introns and two NLS signals flanking the *Cas9*. Cas9i has been reported to dramatically improve the editing efficiency already in the primary T1 transformants. However, according to Grützner et al., none of the primary transformants obtained with a *Cas9* lacking introns driven under *pRPS5a* promoter displayed a knockout mutant phenotype, whereas between 70% and 100% of the transformants generated with the Cas9i displayed mutant phenotypes (Grützner et al., 2021). The inefficiency of non-modified Cas9 in this report was not matching with our data, making us wonder what improvement in efficiency of Cas9i could be expected in our system with a better-working, unmodified Cas9.

In addition to *Cas9i*, we also replaced the FastRed cassette with FastGreen and a newly generated FastCyan to produce a set of destination vectors containing *Cas9i* driven either under *pEC1.2* or *PcUBi4-2* and combined with FastRed, FastGreen or FastCyan fluorescent seed selection markers (Fig. 4B, Table 1 and Supplemental Fig. 1). To test the efficiency of these newly generated plasmids, we chose *SGN3 (SCHENGEN 3)* as a target gene. *SGN3* encodes a receptor-like kinase with an important function in maintaining the integrity of Casparian strips, lignin-based barrier in the root endodermis. In the absence of SGN3, discontinuous patches of lignin can be easily observed using Basic Fuchsin staining (Pfister et al., 2014; Doblas et al., 2017; Fujita et al., 2020). For comparison, we introduced three gRNAs to target *SGN3* into the vectors containing *Cas9i* and *Cas9* lacking the introns (Fig. 4C). In both cases, *Cas9* or *Cas9i* were driven either under *PcUBi4-2* or *pEC1.2* promoters. In T1 generation, we assessed the number of *sgn3* knockout phenotype appearing in different lines. We observed that the highest number of knockout phenotypes (32/51, 62.7%) were displayed by the T1 line generated using intron-optimized *Cas9i* driven under *pEC1.2* promoter. In comparison, the same promoter driving regular, intron-less Cas9, generated 30% (15/50) knockout phenotype displaying individuals (Fig. 4D–E). Although *Cas9i* driven under ubiquitous *PcUBi4-2* promoter generated significantly lower number of knockouts in T1 generation (5/50, 10%), it was still higher compared to the same promoter driving regular *Cas9* (3/50, 6%) (Fig. 4E). To check the inheritance of the knockout phenotypes, we generated T2 lines and checked two lines from each construct, *pEC1.2::Cas9* and *pEC1.2::Cas9i*. In all four lines we could confirm homozygous *sgn3* phenotype by selecting dark, Cas9-free, seeds (Fig. 4F). These results show that Cas9i indeed represents a significant improvement over intron-less variants, even when compared in a system in which intron-less Cas9 shows a reasonable efficiency. Our data indicates that the *pEC1::Cas9i* construct reported here is a highly efficient vector to efficiently produce biallelic mutants already in T1 generation.

## DISCUSSION

In this study, we developed and tested a two-color based multiplex CRISPR/Cas9 toolbox that consists of Golden Gate- and Gateway-compatible vectors, which will be made available at Addgene. The vectors allow to assemble up to eight (sixteen in case of co-dipping) gRNAs. The T-DNA vectors we developed contain either FastRed, FastGreen or FastCyan cassette which: 1) simplifies the cloning strategy by avoiding the generation of very large constructs; 2) simplifies the screening method in the first generation (T1) where only fluorescent seeds are selected, as well as in second generation (T2) where Cas9 presence is counter-selected by picking non-fluorescent seeds and which are then checked for the desired editing events; 3) provides ability to easily screen for different, higher order mutants in the same generation. To demonstrate multiplexing, we cloned three independent gRNAs in each vector simultaneously, co-transformed them together into Arabidopsis, an approach that has not yet been explored. The T-DNA vectors contain either *pEC1.2* or *PcUBi4-2* promoter to drive Cas9, intron-optimized Cas9, SaCas9 or AsCpf1 and fluorescent seed selection cassette of choice – FastRed, FastGreen or FastCyan. Moreover, most of the vectors we generated can be easily modified to be able to introduce any promoter, Cas gene or selection cassette of interest.

We believe the toolbox presented here will be very useful in plant research and plant synthetic biology, due to its streamlined, easy-to-use and efficient cloning and selection system. Moreover, its modularity and flexibility will allow researcher to easily build-on and improve the system in the future.

## MATERIALS AND METHODS

### Plant growth and transformation

In all experiments Arabidopsis thaliana Columbia (Col-0)ecotype was used. The seedlings were germinated on solid half-strength Murashige and Skoog (MS) medium without addition of sucrose. *CUC1 (At3g15170), CUC2 (At5g53950)* and *SGN3 (AT4G20140)* genes were chosen as targets for CRISPR/Cas9 targeting. The seeds of T1 and T2 generations were surface sterilized, sown on plates, incubated for 2 days at 4°C for stratification, and grown vertically in growth chambers at 22°C, under continuous light. The phenotypic analyses were performed on 6-day-old seedlings. For Agrobacterium-mediated transformation, siliques of flowering plants were removed and a solution of resuspended Agrobacterium cells carrying corresponding CRISPR constructs with sucrose and SILWETT (5% of sucrose and 0.06 % Silwet L-77) was directly applied to flower buds by pipetting. In case of co-transformation, FastRed and FastGreen vectors were transformed separately into Agrobacterium and grown overnight in 5ml cultures at 28°C. The cultures were centrifuged for 10 min at 4000 rpm, the pellets resuspended in sucrose and Silwet L-77 solution. The resuspended FastRed and FastGreen pellets were mixed in equal amount to make a cocktail for transforming both constructs at same time.

### Generation of CRISPR/Cas9 vectors

The primers used to generate all vectors are indicated in Supplementary Table 1. pChimera (Addgene ID 61432) (Fauser et al., 2014) was used as a template to generate *pRU41, pRU43, pRU45* and *pRU47* vectors. The *pRU42, pRU44, pRU46* and *pRU48* were generated by replacing the *pU6* promoter with *pU3*. The corresponding *BsaI* sites were introduced to generate compatible overhangs in all the entry clones as shown in Figure 1. The intermediate vectors *pSF463, pSF278, pSF464, pSF279, pSF280, pSF325* were generated by introducing the corresponding BsaI sites into *pDONR221* containing *ccdB* cassette and flanking attL1 and attL2 recombination sites ready for single fragment Gateway cloning. The final T-DNA vectors were generated using *pDe-Cas9* as template (Addgene ID 61433) (Fauser et al., 2014). The FastRed, FastGreen and FastCyan selection cassettes were amplified from *pFRm43GW* (Addgene ID 133748), *pFG7m34GW* (Addgene ID 133747) (Wang et al., 2020) and *UBQ::NLS-mTurquoise2* (Emonet et al., 2021) vectors and introduced into *pDe-Cas9* vector in place of PPT selection using *HindIII* restriction sites. Different *Cas9* and *Cpf1* variants, as well as *Cas9i* were amplified from vectors *pDe-SaCas9* (Steinert et al., 2015), *pYPQ220* (Addgene ID 86208) (Tang et al., 2017) and *pAGM47523* (Addgene ID 153221) (Grützner et al., 2021) and introduced by replacing *Cas9* in *pDe-Cas9* vector. *pEC1.2* was amplified from *pHEE401E* (Addgene ID 71287) (Wang et al., 2015) vector and introduced into different vectors by replacing the *PcUBi4-2* promoter vector using *KpnI* restriction sites. All vectors generated in this study are shown in Table 1, Supplemental Figure 1 and Supplemental Tables 1 and 2 and are deposited at Addgene plasmid repository. Primers used for modifying the vectors are indicated in Supplemental Table 3. The primers for introducing the required gRNAs into entry vectors are indicated in Supplementary Table 4 and the detailed procedure of generating T-DNA vectors carrying the gRNAs is described in Supplemental Materials and Methods.

### Screening of CRISPR mutants

Fluorescent seeds of T1 plants carrying FastGreen, FastRed and FastCyan cassettes, as well as dark (non-fluorescent) seeds of T2 lines were screened using Leica MZ16FA Fluorescence Stereomicroscope. The filters used for different colours are as follows: DSR (LEICA 10447227) for FastRed; GFP3 (LEICA 10447217) for FastGreen; CFP (LEICA 10447409) for FastCyan; GFP2 (LEICA 10447221) for yellow seeds carrying both FastRed and FastGreen constructs. Genomic DNA of transgenic CRISPR T1 and non-transgenic T2 plants was extracted using modified CTAB method. The leaves and flowers were used for DNA extraction. The plant material was crashed using pipette tips directly in 100μl CTAB buffer and incubated for 40 min at 65C. 100μl of chloroform/isoamyl alcohol (16/1 ratio) was added, mixed by inverting and centrifuged for 5min at max speed. The upper phase was collected, mixed with 50μl of isopropanol and incubated overnight. Next day, the samples were centrifuged for 10 min at maximum speed. The liquid was discarded and, after drying, the pellet was resuspended in 50 μl of water. Primers used for genotyping and sequencing are indicated in Supplementary Tables 5 and 6.

### Lignin staining and confocal microscopy

ClearSee‐adapted Basic Fuchsin staining for lignin was performed as described earlier (Ursache et al., 2018). Confocal pictures of Basic Fuchsin stained sgn3 mutant roots were obtained using Zeiss LSM 880 confocal microscope. The excitation and emission spectra for Basic Fuchsin are 561 nm and 570–650 nm accordingly.

## ACKNOWLEDGMENTS

We would like to thank Professor Holger Puchta for providing us with pChimera (Addgene ID 61432), pDe-Cas9 (Addgene ID 61433) and pDe-SaCas9 plasmids. This work was funded by European Molecular Biology Organisation (EMBO ALTF 1046-2015) to R.U., Overseas Research Fellowship form JSPS to S.F and Swiss National Science Foundation (SNSF) grants 310030E_176090, 31003A_156261 and 310030B_176399 to N.G.

## Supplementary Materials and Methods CRISPR/Cas9 Cloning Protocol

### 1. Primer design

Pick your 20 nucleotide protospacer sequence and order desalted oligos:

- **For pRU41 (pU6-gRNA1) vector:**

Forward primer: 5′-**ATTG** + protospacer
Reverse primer: 5′-**AAAC** + rev-com protospacer
- **For pRU42 (pU3-gRNA2) vector:**

Forward primer: 5′-**GTCA** + protospacer
Reverse primer: 5′-**AAAC** + rev-com protospacer
- **For pRU43 (pU6-gRNA3) vector:**

Forward primer: 5′-**ATTG** + protospacer
Reverse primer: 5′-**AAAC** + rev-com protospacer
- **For pRU44 (pU3-gRNA4) vector:**

Forward primer: 5′-**GTCA** + protospacer
Reverse primer: 5′-**AAAC** + rev-com protospacer
- **For pRU45 (pU6-gRNA5) vector:**

Forward primer: 5′-**ATTG** + protospacer
Reverse primer: 5′-**AAAC** + rev-com protospacer
- **For pRU46 (pU3-gRNA6) vector:**

Forward primer: 5′-**GTCA** + protospacer
Reverse primer: 5′-**AAAC** + rev-com protospacer
- **For pRU47 (pU6-gRNA7) vector:**

Forward primer: 5′-**ATTG** + protospacer
Reverse primer: 5′-**AAAC** + rev-com protospacer
- **For pRU48 (pU3-gRNA8) vector:**

Forward primer: 5′-**GTCA** + protospacer
Reverse primer: 5′-**AAAC** + rev-com protospacer

### 2. Cloning gRNAs into pRU41-48 using Oligo annealing

1 μl of each oligo (100 μM) + 48 μl H20
Incubate for 5 min at 95°C (thermocycler, no cooling at the end!)
Cooling at room temperature for 20 min

- **Digest entry vector:**

5 μl of corresponding pRU41-pRU48 entry vector
1 μl of FastDigest Buffer (ThermoScientific)
1 μl FastDigest BbsI (Bpi) enzyme (Thermo Fisher)
adjust water to 10 μl final volume and incubate for >1 h at 37°C (overnight is optimal)
Gel extract the digested vector and adjust the concentration to 5 ng/μl
- **Ligation**

μl of corresponding digested pRU41-48 entry vector
3 μl of annealed oligos
1,5 μl of T4 Ligase (ThermoScientific)
2 μl T4 buffer
1.5 μl H_2_O
Incubate for at least 1h at 22 °C or room temperature
Transform everything in DH5α, plate on LB plates supplied with Ampicillin
- **Colony-PCR**

Test 4 colonies (efficiency >70%) using oRU385 + gRNA reverse oligo
- **Miniprep**

Sequence using oRU385 primer and adjust plasmid concentrations to 100 ng/μl

### 3. Golden Gate Assembly

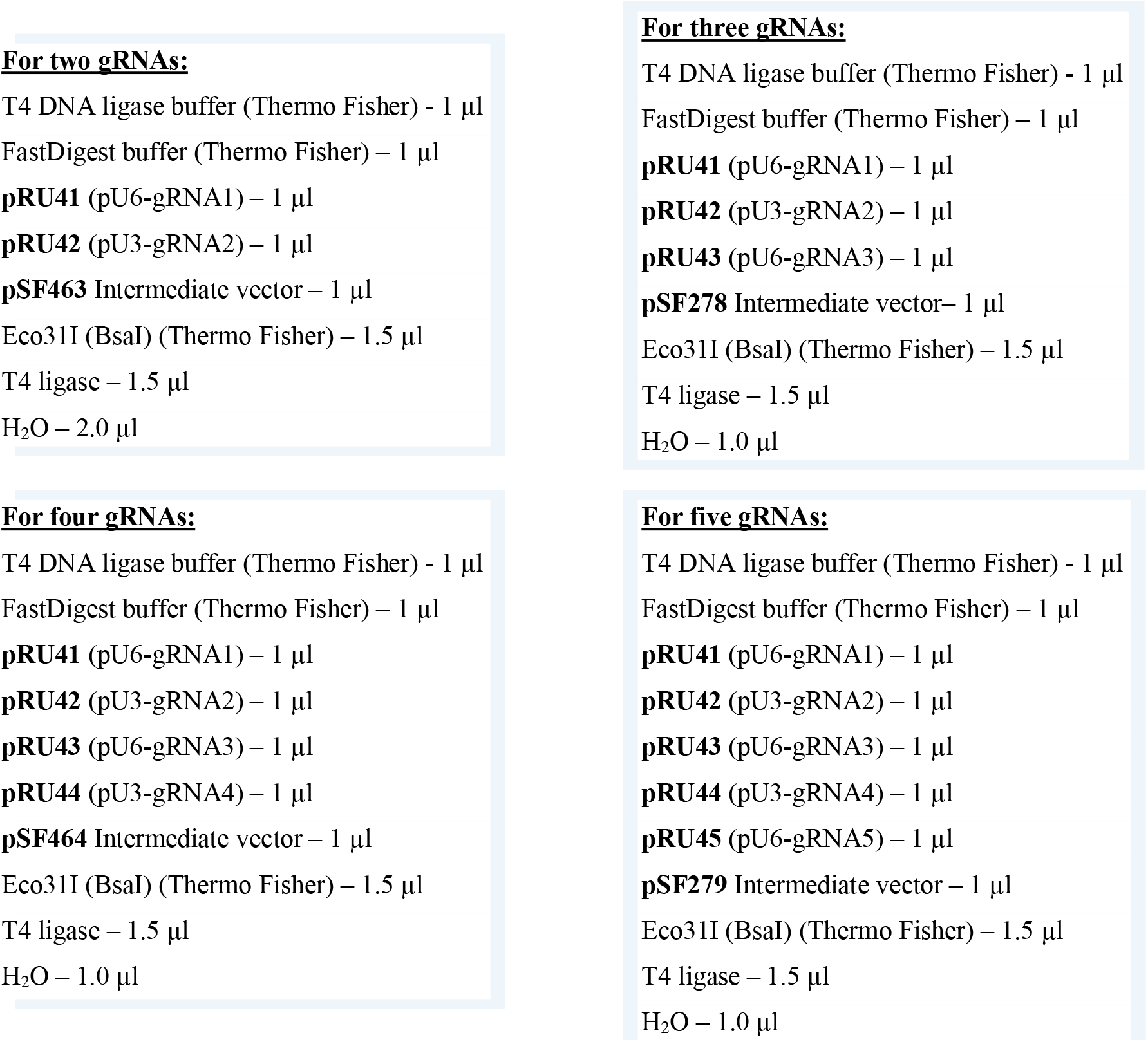
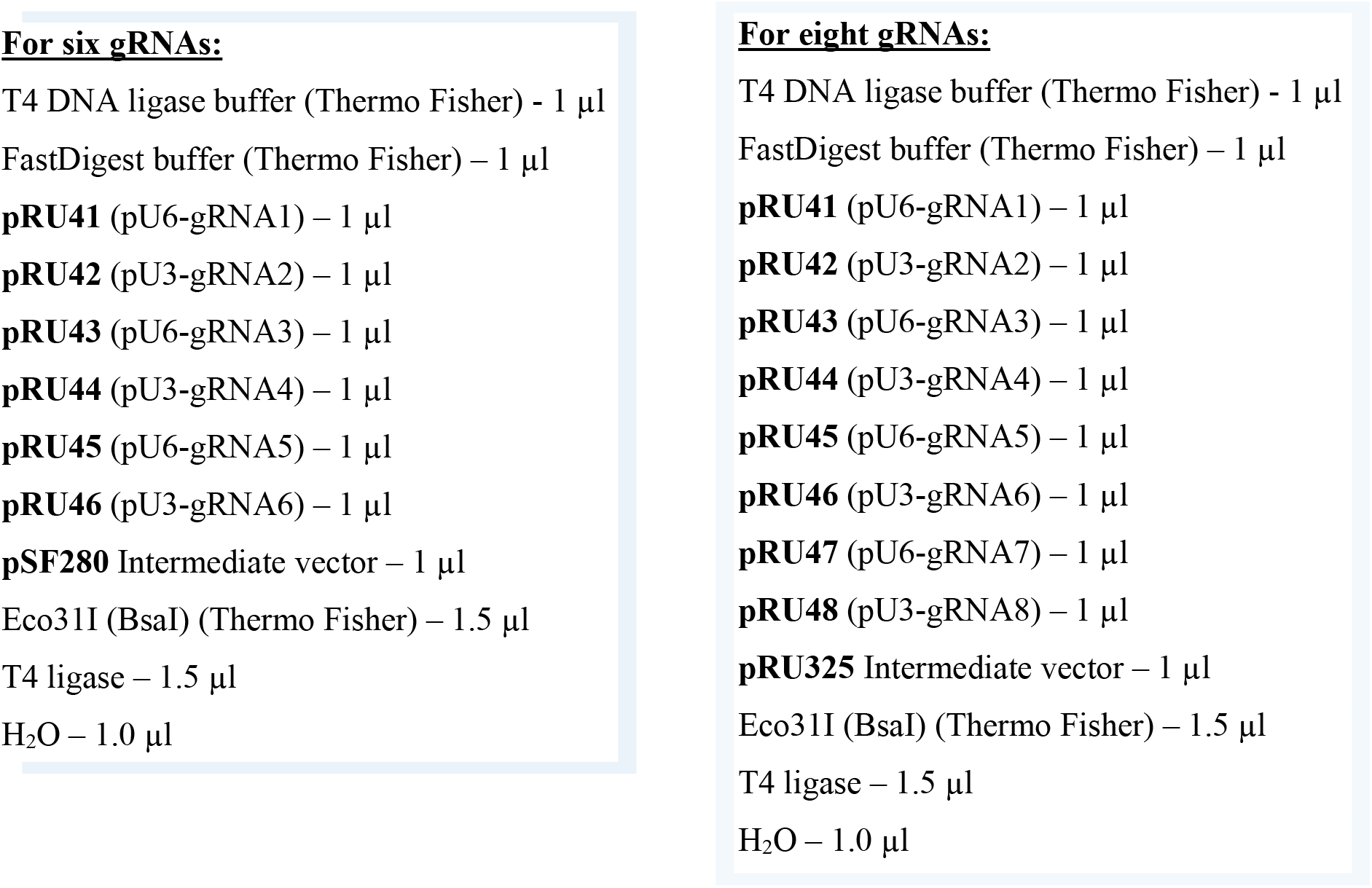

#### Run Golden Gate program in a thermocycler as follows

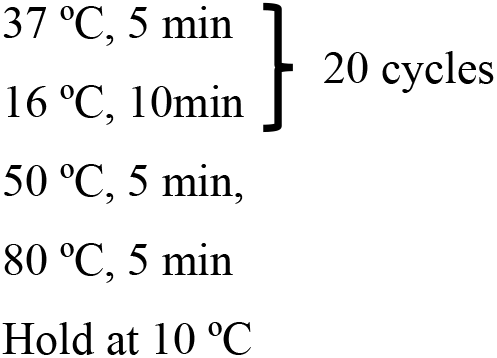

- Transform everything into E.coli DH5α cells and plate transformed cells on LB/Kan plates.
- Check 2-4 colonies, miniprep, sequence using M13 and M13 reverse primers. To check the gRNA’s in the middle, gRNA primers can be used as colony PCR and sequencing primers.

### 4. Final Single Fragment Gateway LR Reaction

2 μl of your intermediate vector with gRNAs assembled (adjusted to 50 ng/μl)
3 μl of the final Cas9 vector (adjusted to 50 ng/μl)
4 μl TE buffer, pH 8
1 μl LR clonase II

- Incubate overnight at room temperature
- Proteinase K treatment: add 1 μl and incubate for 10 min at 37°C
- Transform everything in DH5α and plate everything on LB plates with Spectinomycin
- Miniprep, all colonies should be positive, inoculate 1-2 colonies
- Sequence using oRU906, oRU908 primers or gRNAs as primers

#### Primer Sequences

oRU906: GAGTCTATGATCAAGTAATTATGC
oRU908: GCTTGCATGCCTGCAGGTCGACTCT
oRU385: CAACGCGTTGGGAGCTCTCCCATATG

## Supplemental Tables

**Supplemental Table 1.**
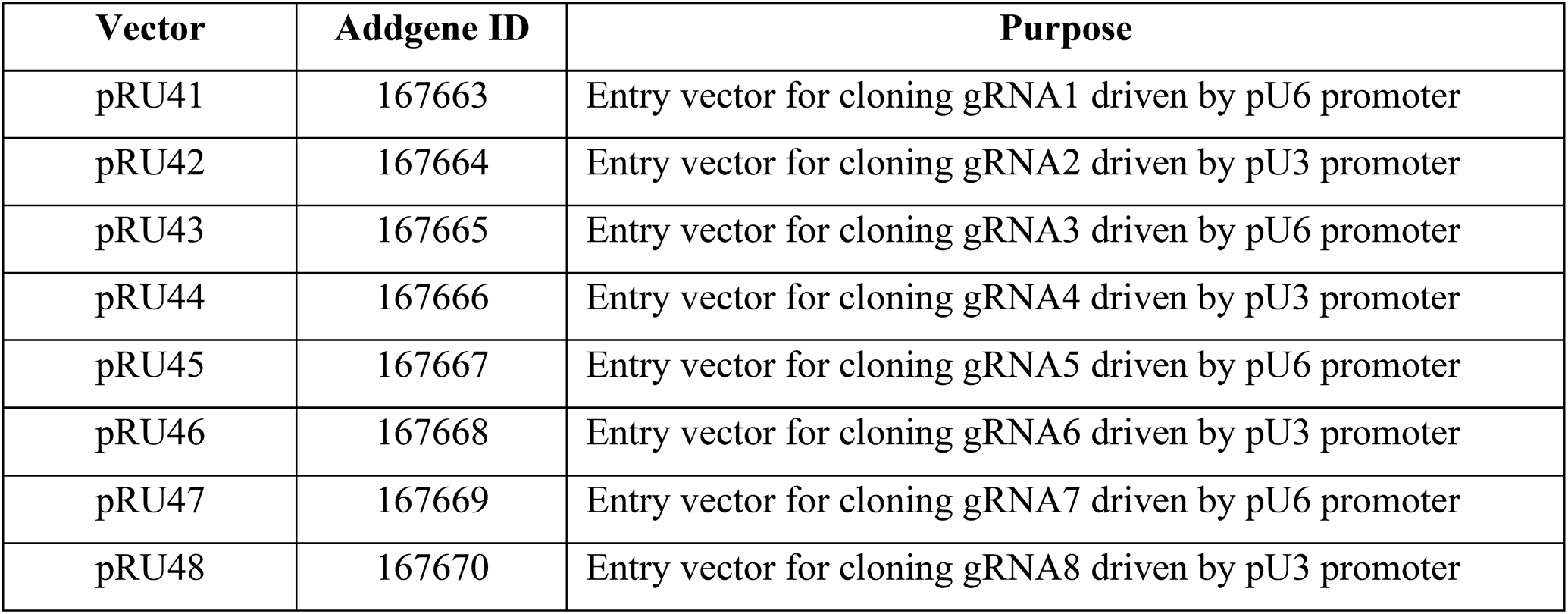
1st level intermediate vectors.

**Supplemental Table 2.**
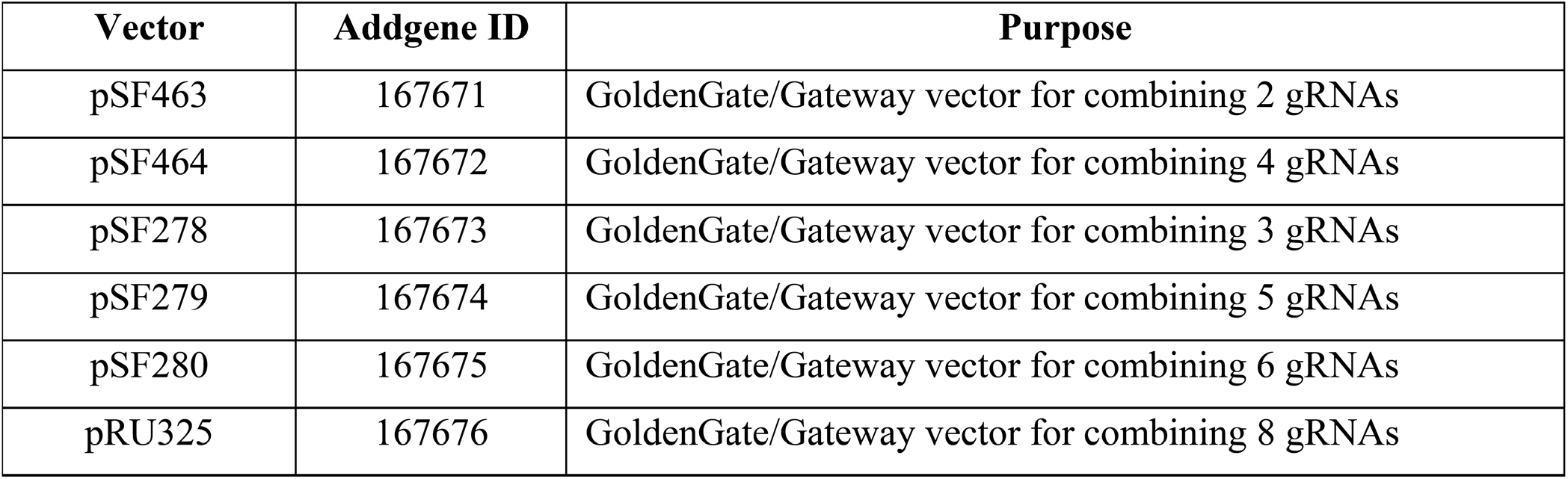
2nd level intermediate vectors

| Vector | Addgene ID | Purpose |
| --- | --- | --- |
| pSF463 | 167671 | GoldenGate/Gateway vector for combining 2 gRNAs |
| pSF464 | 167672 | GoldenGate/Gateway vector for combining 4 gRNAs |
| pSF278 | 167673 | GoldenGate/Gateway vector for combining 3 gRNAs |
| pSF279 | 167674 | GoldenGate/Gateway vector for combining 5 gRNAs |
| pSF280 | 167675 | GoldenGate/Gateway vector for combining 6 gRNAs |
| pRU325 | 167676 | GoldenGate/Gateway vector for combining 8 gRNAs |

**Supplemental Table 3.**
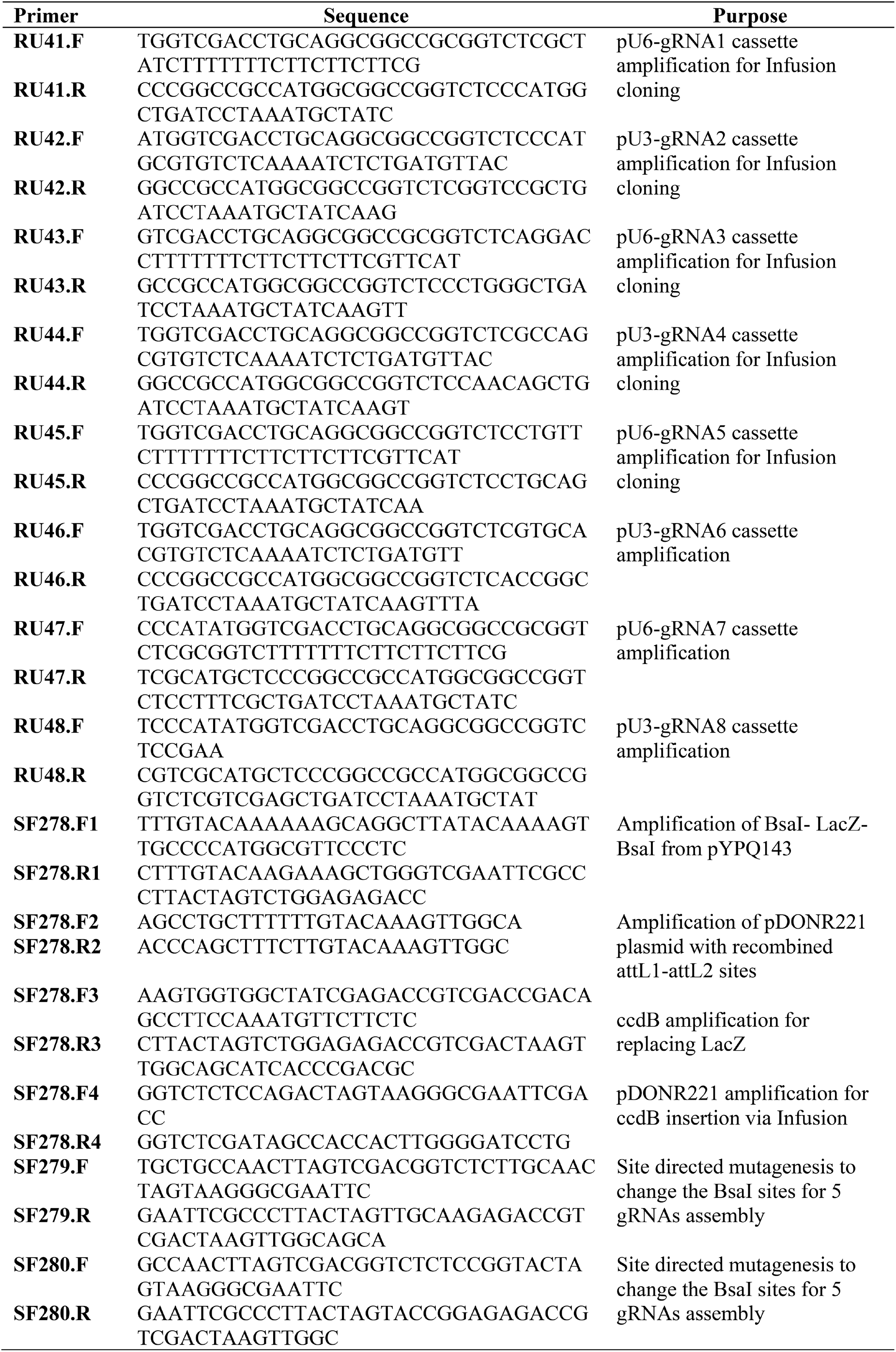
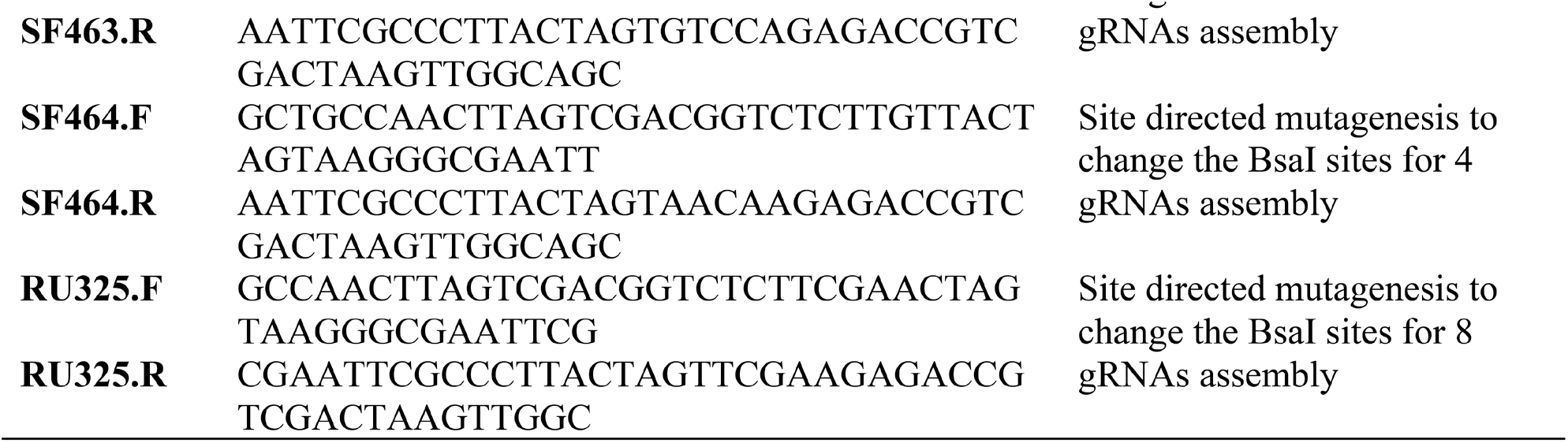
Primers for generating the entry and intermediate vectors

**Supplemental Table 4.**
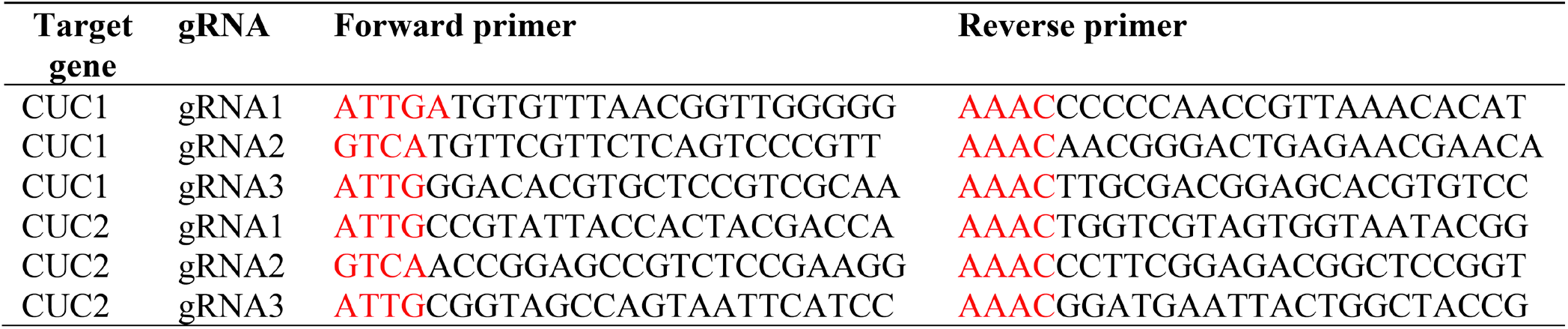
Primers for introducing the gRNAs using oligo anealing technique. Red color marks the overhangs required for ligation

**Supplemental Table 5.**
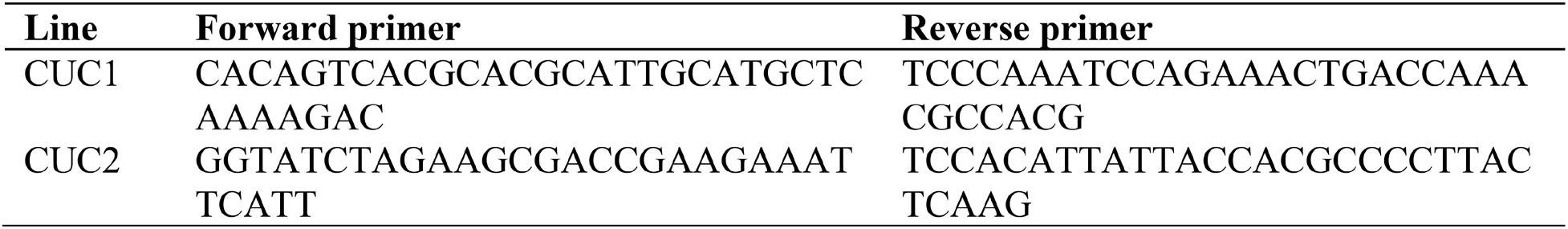
Primers used for PCR-based genotyping

**Supplemental Table 6.**
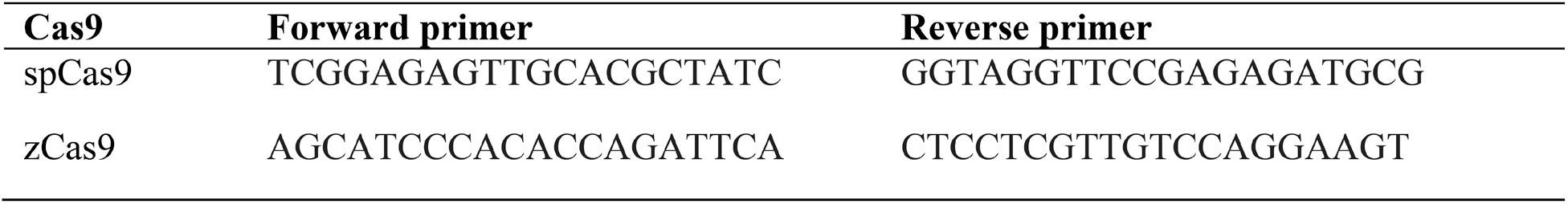
Primers used for Cas9 amplification

**Supplemental Figure S1.**
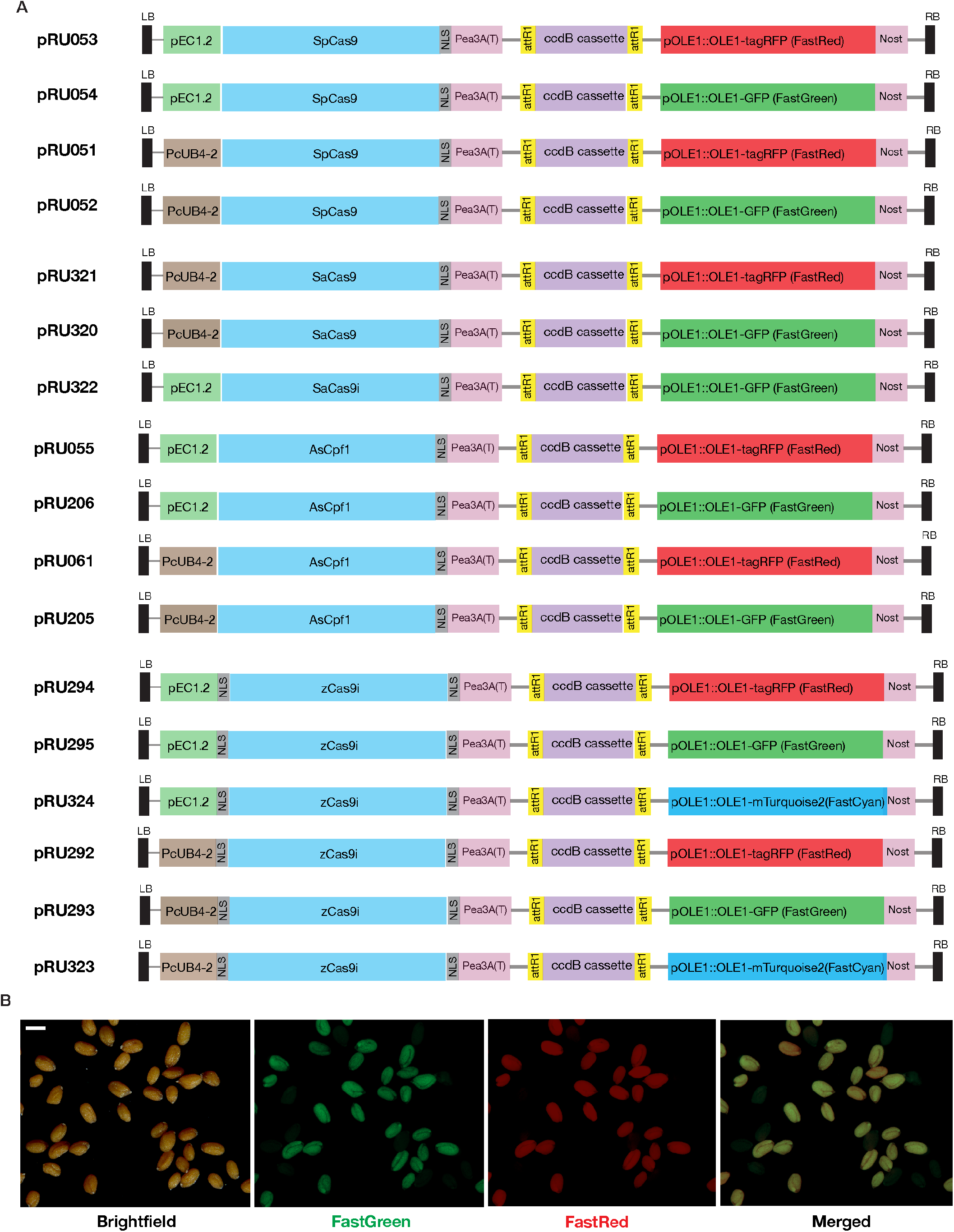
A, Schematics showing the collection of final T-DNA vectors developed in this study. B, Example of dominant presence of double-color fluorescent seeds in T2 generation. Scale bar = 100 μm.

